# Degradable endovascular neural interface for minimally invasive neural recording and stimulation

**DOI:** 10.1101/2021.03.24.436737

**Authors:** Adele Fanelli, Laura Ferlauto, Elodie Geneviève Zollinger, Olivier Brina, Philippe Reymond, Paolo Machi, Diego Ghezzi

**Affiliations:** Medtronic Chair in Neuroengineering, Center for Neuroprosthetics and Institute of Bioengineering, School of Engineering, École polytechnique fédérale de Lausanne, Chemin des Mines 9, 1202 Geneva, Switzerland; Division of of Neuroradiology, Geneva University Hospital, Geneva, Switzerland

**Keywords:** neural interfaces, transient bioelectronics, stents, endovascular devices, biodegradable polymers

## Abstract

Neural recording and stimulation have been widely used to mitigate traumatic injuries, neurodegenerative diseases or mental disorders. Most neural interfaces commonly require invasive surgery, potentially entailing both transient and permanent complications. A promising strategy designed to overcome these risks involves exploiting the cerebrovascular system as an access route to the neural tissue. Here we present a novel endovascular neural interface for neural recording and stimulation, fully polymeric and degradable. This concept might allow for better integration of the device in the body, reduced inflammatory reaction, the possibility of replacing the implant after degradation, and avoiding removal surgeries. The vasculature’s strategic distribution and the use of soft polymers for the device’s fabrication will permit targeting both the brain vasculature and the peripheral system. Therefore, this novel endovascular neural interface will broaden the range of applications from neurological diseases and mental disorders to bioelectronics medicine.

## 1. Introduction

Neural recording and stimulation opened the doors to a deeper understanding of the brain and, at the same time, led to the development of treatments for heavily-impairing conditions following traumatic injuries, neurodegenerative diseases or mental disorders.^[1]^ Although the potential impact of neurotechnology is enormous, its clinical use nowadays is still limited. Many systems require invasive surgery, with related high risks that not always overcome the benefits for the patients. A big step in this direction is developing endovascular neural interfaces since endovascular procedures are considerably less invasive, are routinely performed and allow for short patient recovery times.^[2]^ Also, endovascular neural interfaces do not involve craniotomies, reducing the risks associated with the surgical procedure (e.g., susceptibility to seizures) and, therefore, increasing the possibility of patients undergoing these innovative treatments.

Endovascular neural interfaces interact with the surrounding neural tissue from within blood vessels. Devices developed so far consisted mainly of wires or catheters used as electrodes. Even though wires and smart catheters opened the doors to endovascular neural recording and stimulation, their main limitations were the short-term application and the low spatial resolution. The use of a stent-electrode array (e.g. Stentrode™) overcame these issues^[2]^, and advantages of this technology have been recently demonstrated in preclinical and clinical trials.^[3–6]^ Seventeen potential medical targets have been identified where intravascular neuromodulation could replace current invasive deep brain stimulation protocols, among which five with an intraluminal diameter larger than 1 mm.^[7]^ By accessing the internal cerebral vein or the anterior communicating artery, it would be possible to perform minimally invasive neuromodulation for a wide range of neurological disorders such as Parkinson’s, Alzheimer’s, essential tremor, depression and obsessive-compulsive disorders.^[7]^ Also, an endovascular approach allows for minimally invasive neural recording for advanced brain-computer-interfaces.^[3–5]^ Furthermore, endovascular devices could be as well applied to the peripheral system for pain relief, motor deficits, bladder control and muscle’s stimulation, among others. However, the main limitation of stent-electrode arrays is the metallic and conductive substrate for electrodes that makes insulation critical and shortcuts frequent.^[3]^ Also, the use of a metallic stent could induce a strong inflammatory response. A severe foreign body reaction response can affect the performances of a neural interface.^[8–11]^ For stent technology, this limitation already led to the development of bioresorbable devices, allowing a better interaction with the tissue and placement in mechanically solicited areas.^[12]^

These present limitations motivated us to develop a degradable stentrode technology based on polymeric materials and microfabrication techniques. Reducing the inflammatory response will better preserve the neural interface’s functionality, allowing extended durability and better signal quality.^[9,13–16]^ Moreover, whereas in the case of device failure it will not be easy to replace a metallic stentrode, if not impossible at all, having a degradable neural interface will leave the possibility to further implant and make therapy adjustments, even if complications occur in the mid-long-term. Transient bioelectronics already introduced degradable materials in neurotechnology^[17]^. Unfortunately, these devices have a limited lifetime, from days to a few weeks, due to fast degradation rates of the dissolvable metals usually used for the conductive elements. This limited lifetime prevents their use for mid- and long-term applications.^[18,19]^ Our strategy to overcome this problem is a fully polymeric neural interface, using a biodegradable polymer as a substrate and encapsulation and a conductive polymer as the active component.

In summary, in this work, we introduce a fully polymeric degradable endovascular neural interface combining the advantages of transient bioelectronics and stent technology in a single device.

## 2. Results and Discussion

### 2.1. Device’s manufacturing

The degradable endovascular neural interface is a stent-inspired structure, using the vasculature as a minimally invasive access route to the neural tissue. The device’s scaffold is made out of poly-ε-caprolactone (PCL): an FDA-approved synthetic polyester widely used for drug delivery systems, tissue engineering and medical devices.^[20]^ Among the other synthetic biodegradable polymers, PCL has a longer degradation time, from several months to a few years, allowing it to be an insulating and stable scaffold for mid- and long-term applications.^[20]^ Degradable metallic electrodes are made out of poly(3,4-ethylenedioxythiophene):polystyrene sulfonate (PEDOT:PSS), a conductive biocompatible polymer commonly used in organic photovoltaic cells, and often used as a coating for metallic electrodes of neural interfaces.^[21–23]^ A novelty of this device is the absence of metals in electrodes and tracks, making the device fully polymeric in all components. PEDOT:PSS is here employed as the only active component, doped with ethylene glycol (EG) for better conductivity, and (3-glycidyloxypropyl)trimethoxysilane (GOPS) for better endurance to aqueous environments.^[23,24]^

The device contains four polymeric electrodes, with a diameter of 600 μm and a geometrical surface area (GSA) of 0.0019 mm^2^ (electrode’s opening of 500 μm). Polymeric feedlines are 200-μm wide. The electrodes are distributed over the device’s surface to cover an area of approximately 7.6 × 3.6 mm^2^ (**Figure 1a**). A V-shaped area at the device’s tip is used to smoothen the device’s insertion into the catheter. The device’s length is 44 mm, and the largest width is 7.6 mm. The degradable endovascular neural interface is designed to expose polymeric electrodes around a cylindrical surface (e.g., vessel wall), as representatively shown in a 2-mm channel within a poly(methyl methacrylate) (PMMA) block (Figure 1b). Fabrication of the device using replica moulding proved to be simple, fast, and effective. Two moulds have been manufactured in SU8 (35-μm thick): one with the engraved pattern of the tracks (mould for encapsulation in Figure 1c) and the other with the protruding pattern of electrodes and traces (mould for substrate in Figure 1c). After covering the two moulds with a PSS release layer, the PCL solution was spin-coated on both. After releasing the substrate layer, a PEDOT:PSS-based solution was dispensed along the engraved electrodes, feedlines and pads. For each track, approximately 2-μl of conductive polymer would completely fill the pattern. This amount would correspond to a film’s thickness of approximately 1-2 μm, with thickness decreasing from the channel wall to the centre (Figure 1d). After cross-linking of the conductive polymer, thin stainless-steel wires (114-μm diameter) were attached to the pads using a conductive paste. After alignment of the encapsulation layer, with electrodes openings opportunely laser cut, to the substrate, heating the two layers at 60 °C allowed the quick fusion of the two and fixed the wires contacts in between. Finally, the device outline and its characteristic diamond-like holes were laser cut to allow for easier bending, reduce the amount of material implanted and maintain open sites for molecules exchange from and to the blood flow. A thickness of approximately 77 μm was obtained for the device, limited by the high viscosity of the PCL-based solution and the depth of the engraved electrodes pattern necessary to deposit the PEDOT:PSS-based solution smoothly.

**Figure 1.**
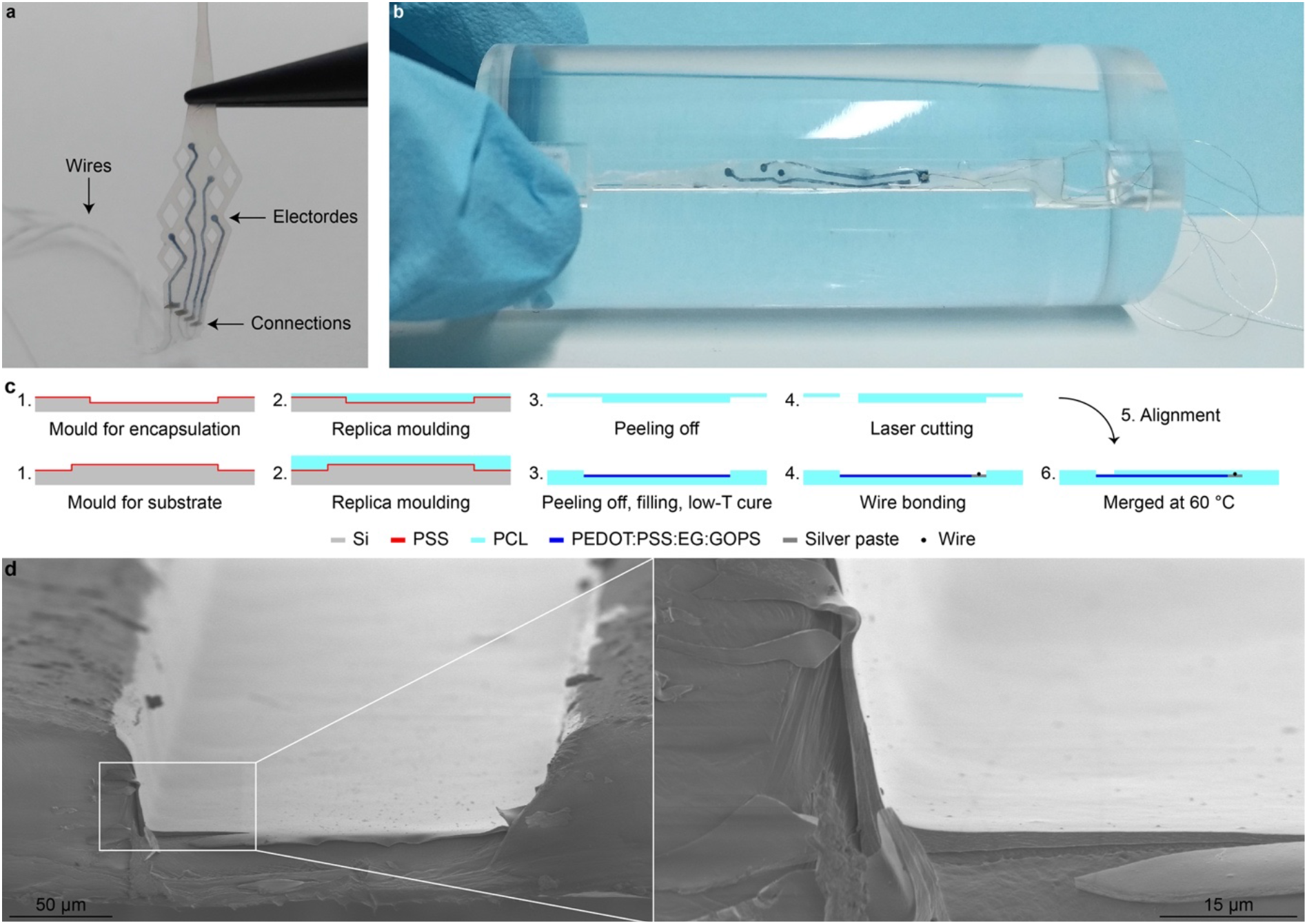
(a) Picture of the degradable endovascular neural interface after fabrication. (b) Picture of the degradable endovascular neural interface deployed in a 2-mm sized channel in a PMMA block. (c) Sketch of the fabrication process. (d) Scanning electron microscope image showing the cross-section of one the feedline filled with the conductive polymer before encapsulation. The conductive polymer layer is visible in the magnification.

### 2.2. Device’s characterisation

After fabrication, we proceeded with the device’s electrochemical characterization. Impedance spectroscopy (**Figure 2a**) performed after fabrication revealed a predominantly resistive behaviour of the electrodes, given by the flatness of curve and the phase angle approaching 0 ° over a wide range of frequencies (10^1^ - 10^5^ Hz).^[25]^ Relevant frequencies in neural recording are included in this range (local field potentials occur at 0.1-300 Hz, and extracellular spikes at 300 - 3000 Hz).^[26]^ At the reference frequency for neural recording (1 kHz), the average (± s.d.) impedance module was 6.7 ± 3.2 kΩ, and the average (± s.d.) impedance phase was −0.56 ± 0.28 ° (*n* = 12 electrodes from *N* = 3 devices; Figure 2c, O condition). Cyclic voltammetry (Figure 2b), performed at a 50 mV s^−1^ scan rate within a window from −0.9 to 0.8 V, showed a rectangular shape indicating a prevalent double-layer capacitance and its integral resulted in a mean total charge storage capacity (CSC) of 42 ± 26.89 mC cm^−2^ (*n* = 12 electrodes from *N* = 3 devices; Figure 2d, O condition).

**Figure 2.**
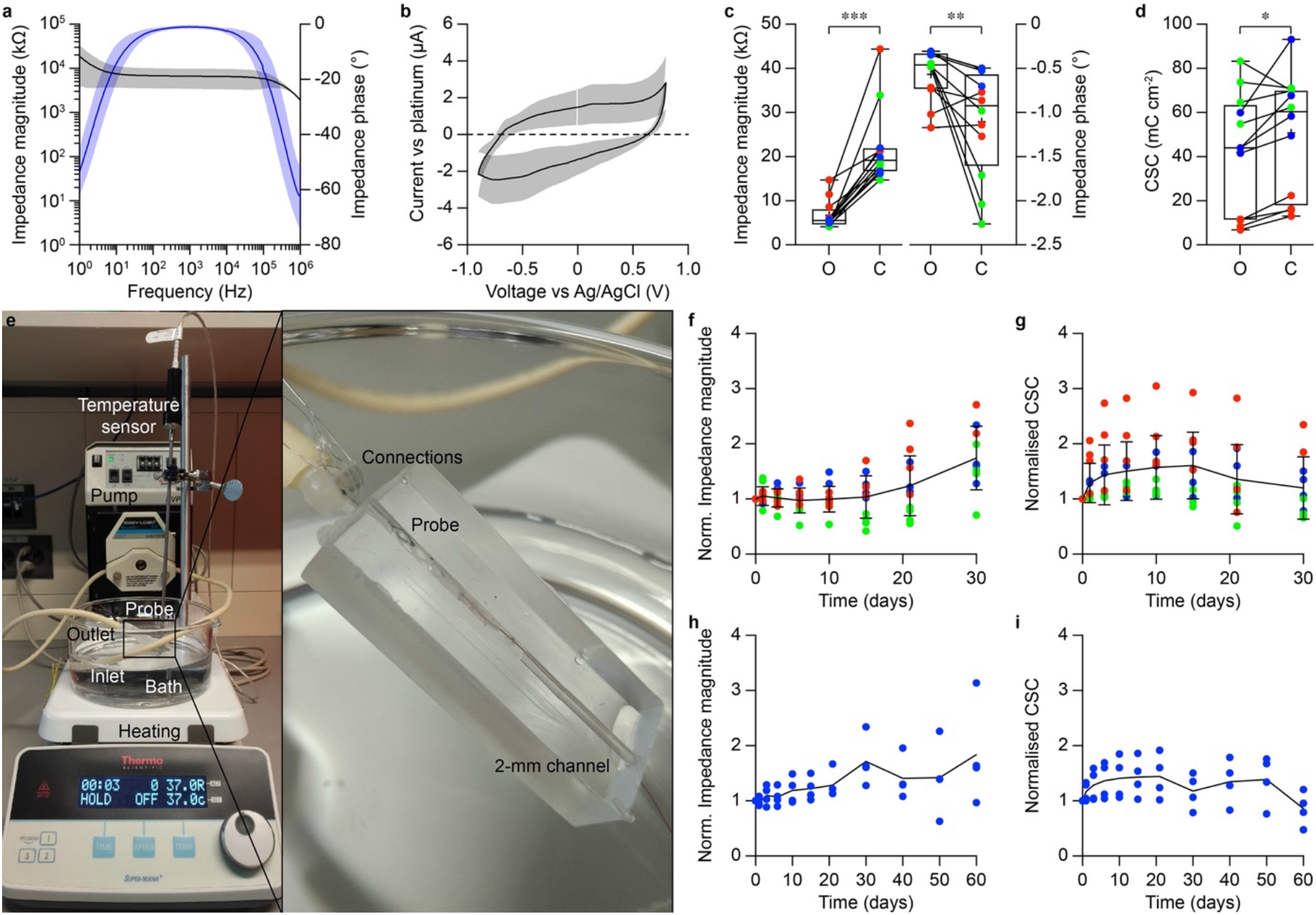
(a) Average (± s.d.) impedance modulus (black, left axis) and impedance phase (blue, right axis) Solid lines are the averages (*n* = 12 electrodes from *N* = 3 devices) and shaded areas are the standard deviations. (b) Average (± s.d.) cyclic voltammetry plot. Solid lines are the averages (*n* = 12 electrodes from *N* = 3 devices) and shaded areas are the standard deviations. (c) Quantification of the impedance magnitude at 1 kHz (left) and impedance phase (right) before (‘O’) and after insertion in 2-mm diameter channel (‘C’); different colours correspond to electrodes from different devices. (d) Quantification of the CSC before (‘O’) and after insertion in 2-mm diameter channel (‘C’); different colours correspond to electrodes from different devices. (e) Picture of the experimental set-up showing: the hot-plate set at 37 °C, the temperature sensor, the beaker containing PBS, and the tubes connected to the peristaltic pump and to the channel where the device is deployed. The magnification shows the device deployed in a 2-mm diameter polydimethylsiloxane channel. (f,g) Normalized impedance magnitude (f) and normalized CSC (g) over 30 days. Different colours correspond to electrodes from different devices. The solid line is the average and bars are the standard deviation. (h,i) Normalized impedance magnitude (h) and normalized CSC (i) over 60 days of one of the 3 devices (blue). The solid line is the average.

Then, the device has been inserted in a 2-mm channel (made out of polydimethylsiloxane) filled with phosphate buffered solution (PBS) at 37 °C and connected to a peristaltic pump, generating a pulsatile flow inside the channel, to investigate the electrochemical behaviour over time in an environment modelling the vasculature (Figure 2e). The pulsatile flow was provided by a peristaltic pump set at 500 rpm corresponding to 106 ml min^−1^ to simulate the cerebral venous flow of Yucatan pigs (108 ml min^−1^), which is the envisioned animal model for further in-vivo validation of the device.^[27]^ Electrodes have been monitored for 30 days, and the impedance module at 1 kHz and the CSC have been used as parameters to assess the electrode’s endurance. First, we observed a change in these values from the after-fabrication condition (O condition) compared to the device inserted in the channel without flow (Figure 2c,d; C condition). The impedance module significantly increased by approximately an order of magnitude (p = 0.0005, two-tailed Wilcoxon matched-pairs test), to a mean (± s.d.) value of 21.9 ± 8.6 kΩ (*n* = 12 electrodes from *N* = 3 devices; Figure 2c). This increase can be explained by the changes in the electrolytic cell’s geometry, owed to the interposition of the channel volume between the device and the reference and counter electrodes which are left in the bath. The impedance phase angle also significantly changed between the two conditions (p = 0.0093, two-tailed Wilcoxon matched-pairs test). The total CSC also significantly increased (p = 0.0254, two-tailed paired t-test) to a mean value of 50.8 ± 27 mC cm^−2^ (*n* = 12 electrodes from *N* = 3 devices; Figure 2d).

Then, the flow was started, and we measured the impedance module at 1 kHz and the total CSC over time, normalized to their respective initial values in the channel (day 0). After 30 days of flow at 37 °C, the impedance at 1 kHz increased to 1.7 times of its original value (*n* = 10 electrodes from *N* = 3 devices; Figure 2f) and the total CSC slightly changed to 1.2 times of its original value (*n* = 10 electrodes from *N* = 3 devices; Figure 2g). The reason why the impedance module increased and, at the same time, the CSC increased as well is not yet clear. The increase in impedance over time might be attributed to two factors: a deterioration of the electrodes or a tightening of the device-wall interface. On the other hand, the increase of the CSC might be explained by the swelling of PEDOT:PSS.^[28–30]^ Furthermore, this swelling would cause electrolyte’s diffusion within the PEDOT:PSS network and its penetration into those areas covered by the encapsulation layer, increasing the effective GSA of the electrode. The electrolyte leakage is detectable at slow scanning rates (50 mV s^−1^) but not at high scanning rates since the pore resistance represents an obstacle limiting the access to the area under the encapsulation.^[25]^ Therefore, cyclic voltammetry would be affected by this phenomenon, while impedance measurements at 1 kHz would not. For a prolonged investigation, one device has been left inside the pulsatile system for additional 30 days: the impedance module reached 1.8 times its original value (*n* = 4, Figure 2h) and the CSC decreased to 0.8 times its original value (*n* = 4, Figure 2i) at day 60. These results suggest that the device is still functional after two months, even if some signs of electrode’s damage appeared (a drop of the CSC and an increase in the impedance variability among the four electrodes).

To further characterize the device for neural stimulation, voltage transients (VTs) have been investigated, and the charge injection capacity (CIC) has been computed. Each electrode of 4 devices has been stimulated with low-current pulses repeated at 150 Hz until stabilization of the interface polarization was reached. Afterwards, the CIC was tested: the injected current has been increased until the most negative polarization value (E_cm_) exceeded the water window of oxidation and reduction potentials (−0.9 V or 0.8 V). All pulses were cathodic first, asymmetric and charge-balanced with a ratio cathodic/anodic equal to 0.2 and 0.05-ms interphase period (**Figure 3**). VTs have a rectangular shape with a significant ohmic drop related to the mostly resistive behaviour of the electrodes, visible as well in the impedance spectroscopy. This VT waveform is maintained for different injected currents within the maximum limit (Figure 3a) and different pulse widths (Figure 3b). Also, the electrode’s equilibrium potential remained constant for stimulations with different pulse widths (from 0.1 to 0.5 ms, steps of 0.1 ms), suggesting no irreversibility in the charge injection. The maximum injectable current has been tested with stimuli of 0.1-ms duration (cathodic phase). The mean (± s.d.) maximum injectable current was found to be 5.2 ± 4.8 mA (*n* = 15 electrodes from *N* = 4 devices) and the respective CIC was 263.6 ± 247.5 mC cm^−2^ (*n* = 15 electrodes from *N* = 4 devices), as reported in Figure 3c. Although these values might change in-vivo, a maximum injectable current of 5.2 mA on average could allow for high-current stimulations without causing relevant damage that might affect the electrodes’ functionality. Currents used in metallic stentrodes fall within the range sustainable by the polymeric electrode presented in this article.^[6]^ The device has then been tested for long-term charge injection by exposing the electrodes to 100’000 pulses at 150 Hz. The amplitude of the cathodic pulse was set to 500 μA, corresponding to 50 nC per phase. The device remained functional (Figure 3d) and the amplitude of the VTs only slightly increased towards the end of the stimulation (Figure 3e), as reflected in the cathodic peaks reached at different moments during the stimulation (p = 0.0648, Friedman test). Electrodes also showed no sign of damage (Figure 3f).

**Figure 3.**
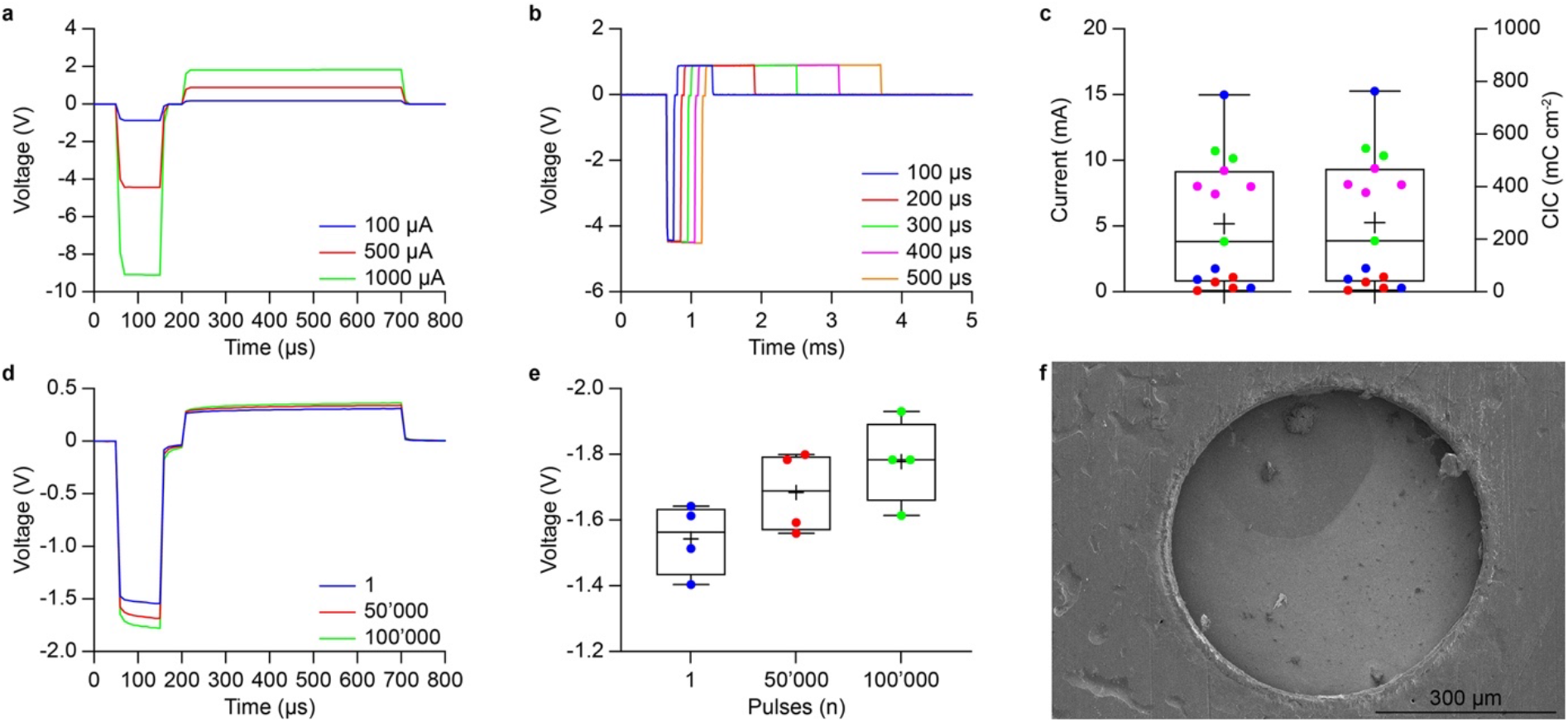
(a,b) Representative VT at different current amplitudes (a) and pulse widths (b). (c) Box plots of the maximum injectable currents (left axis) and CIC (right axis). Different colours correspond to electrodes from different devices. (d) VTs at the beginning (blue), middle (red) and end (green) of prolonged stimulation (100’000 pulses, 500 μA and 150 Hz). (e) Mean (± s.d.) quantification of the cathodic peaks during prolonged stimulation. (f) Scanning electron microscope image of one of the electrodes after being exposed to 100’000 pulses at 500 μA and 150 Hz.

In summary, the device’s electrochemical characterization showed the potential of these fully polymeric electrodes for neural recording and stimulation. The low impedance over a wide frequency range is promising for favourable signal-to-noise ratio in recordings.^[8]^ The high CSC (an index of the electrodes’ charge transfer capabilities) is promising for neurostimulation. It must be noted though that the CSC is an overestimation of the charge transfer potential of the electrodes: in fact when stimulating in-vivo, at high frequencies and short pulses, only the superficial layers of the bulk PEDOT:PSS film will be active.^[8]^ Thus, the CIC was expected to be lower (5-20% of the CSC for metals and oxides).^[25]^ PEDOT:PSS electrodes reported a CIC higher than the total CSC with high variability among electrodes. A similar trend has been reported already in literature for PEDOT-coated PtIr electrodes.^[31]^ A factor contributing to the high CIC is the positive equilibrium voltage reached by the electrodes after a prolonged stimulation at low current pulses to reach stabilization at the electrode-electrolyte interface. This equilibrium voltage was positive for all electrodes (approximately 0.25 V for most electrodes with a few exceptions at 0.15 V), leading most electrodes to a positive E_cm_ value for injected currents below 1500 μA.

### 2.3. Device’s degradation

Next, we tested the device’s degradation profile. An advantage of synthetic degradable polymers is the possibility to tailor their degradation rate. When targeting the vasculature, it is crucial to match the degradation rate to the endothelialisation and remodelling of the blood vessel and avoid chronic recoil and restenosis.^[49]^ A slow degradation is an appealing feature for transient devices since it might allow for better tissue remodelling and reduced chronic trauma at the implantation site.^[32–34]^ On the other hand, the PCL structure will likely last for several years before full degradation: a time that might largely exceeds the functionality of the PEDOT:PSS electrodes and what is needed to allow endothelialisation and tissue remodelling. Therefore, we considered various blends of PCL and poly (D,L-lactic-co-glycolic acid) (PLGA) by mixing 85:15 PLGA with the PCL solution. Mixtures with different ratios of the two compounds have been prepared (i.e. 80:20 and 70:30 PCL:PLGA) and devices have been fabricated and tested for accelerated in-vitro degradation at 37 °C and pH 12 (**Figure 4a,b**). A fast degradation occurred for samples 80:20 and 70:30 since the beginning, while the pristine PCL started degrading around the day 150 (Figure 4a,b). The composition 70:30 degraded to half its initial weight around the day 120. As expected, the degradation of the other mixtures is slower. The acceleration factor at pH 12 is estimated to be 2.5 for PCL (compared to degradation profiles at Ph 7), therefore the composition 70:30 is expected to lose half its initial weight after approximately 10 months.^[32]^ It must be noted that accelerated ageing only evaluated the PCL degradation by hydrolysis, but PCL is also subjected to enzymatic degradation. Specifically, it has been reported that in-vivo PCL undergoes a first degradation by hydrolysis lasting approximately 12 months. Then, after its molecular weight is reduced to about 3,000, it is internalised by the cells (macrophages, giant cells and fibroblasts) and degraded inside them.^[20]^ Therefore, with accelerated in-vitro tests, the device’s degradation time might be overestimated.

**Figure 4.**
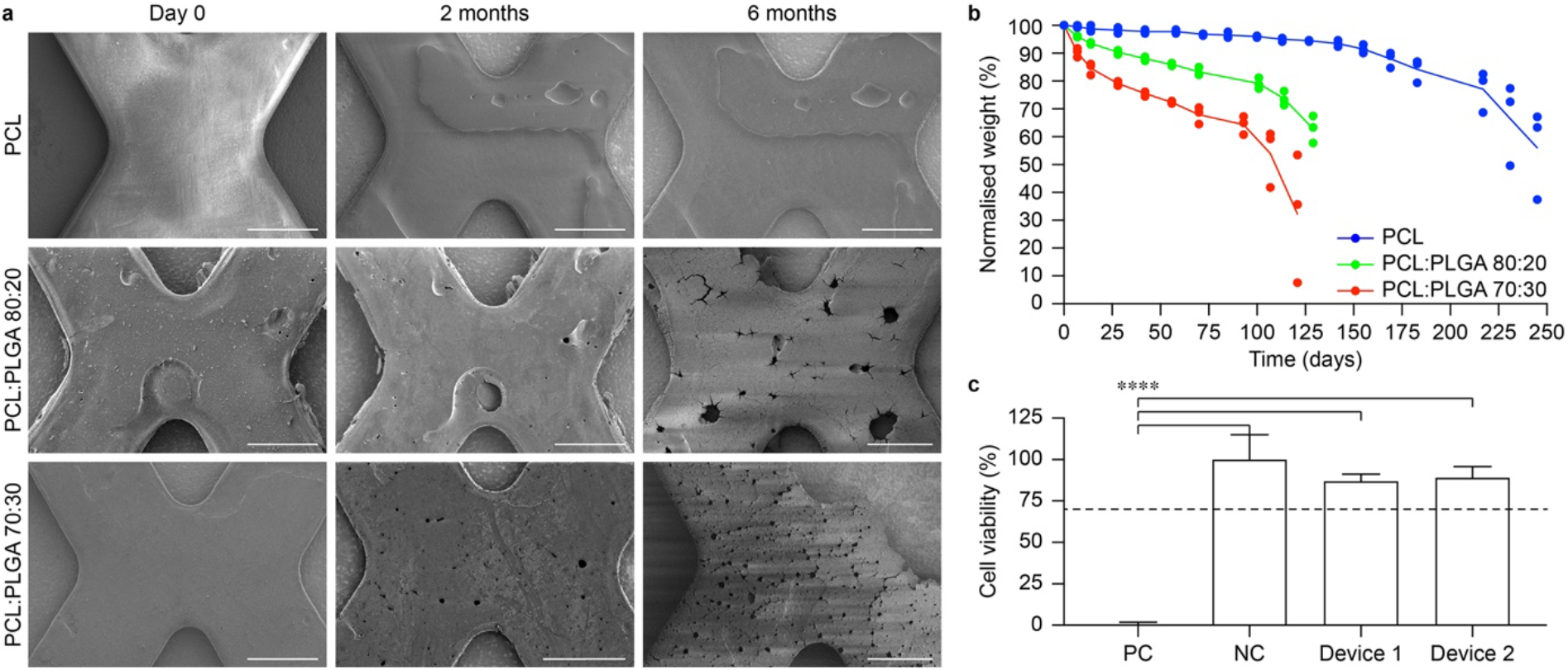
(a) Scanning electron microscope images of PCL, PCL:PLGA 80:20 and PCL:PLGA 70:30 samples during degradation. (b) Normalized weight of PCL (blue), PCL:PLGA 80:20 (green) and PCL:PLGA 70:30 (red) samples during degradation. Solid lines are the averages. (c) Quantification of in-vitro cytotoxicity (mean ± s.d.) for each condition tested: positive control 0 ± 1.81 % (2 samples, 3 replicas per sample); negative control 100 ± 15 % (2 samples, 3 replicas per sample); degradable endovascular neural interfaces 87 ± 4.14 % and 89 ± 6.74 % (2 devices, 3 replicas per device). One-way ANOVA: F = 108.5 and p < 0.0001. Tukey’s multiple comparisons test: positive control vs. negative control p < 0.0001; positive control vs. device 1 p < 0.0001; positive control vs. device 2 p < 0.0001; negative control vs. device 1 p = 0.2891; negative control vs. device 2 p = 0.4234; device 1 vs. device 2 p = 0.9943.

Notably, the device did not show any sign of cytotoxicity (Figure 4c).

### 2.4. Device delivery

After the device’s characterisation and the degradation study, we investigated the device’s compatibility with a standard delivery approach. The shape of the device has been optimised for a smooth insertion inside the delivery catheter thanks to the initial V-shape that allows for a gradual bending and adaptation of the device to the cylindrical constraints of the catheter. The device could be attached to a push wire (0.36-mm diameter) and navigated inside a 5-Fr catheter (1.65-mm outer diameter). When deployed, the device maintains a bent cylindrical shape induced by the insertion and navigation inside the catheter, and, without constraints, tends to expand to restore the planar configuration with which it has been fabricated (**Figure 5a**). This expansion promises good adherence when deployed inside a blood vessel. Also, the navigation inside the catheter and the deployment do not affect the device’s mechanical integrity (Figure 5b,c).

**Figure 5.**
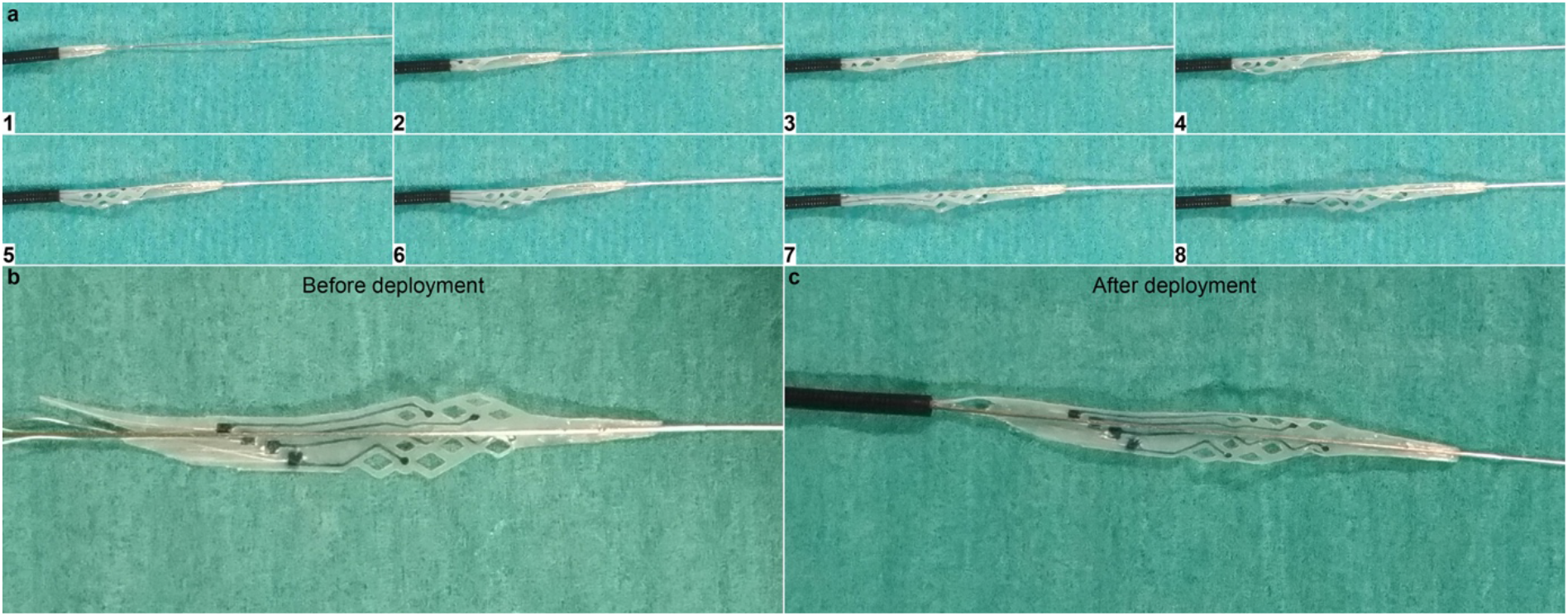
(a) Image sequence of the device deployments using a 5-Fr catheter. (b,c) Comparison of the device before (b) and after (c) deployment.

A micro-computed tomography (micro-CT) scan has been performed on the device deployed inside a 2-mm diameter channel built in a rigid PMMA block to evaluate the device’s apposition once released (**Figure 1b**). Being fabricated in a planar configuration, the device, once deployed, will naturally want to restore its original planar shape and for this will tend to adhere to the vessel walls constraining it in place. The micro-CT scan has been carried out at 10-μm resolution, and both the vertical longitudinal (**Figure 6a**) and transversal (Figure 6b-d) cross-sections of the channel have been analysed. The device’s cross-section inside the channel showed a good device apposition against the channel walls with the vessel lumen almost entirely maintained. Nevertheless, a few gaps between the device and the channel wall were formed. On average (± s.d.), gaps were 98.1-μm long (± 42.4 μm, *n* = 9 for *N* = 3 cross-sections considered), sufficiently far even from moderate gap values (180 μm) reported in the literature that can potentially affect the haemodynamic within the vessel.^[35]^ The lumen left available for the blood to flow delimited by the device was on average (± s.d.) 2.57 mm^2^ (± 0.18 mm^2^, *n* = 9 for *N* = 3 cross-sections considered). Compared to the original lumen area of 3.14 mm^2^, the deployment of the device caused a lumen reduction of 18%. In an ideal scenario of perfect device apposition, the lumen reduction would be 15% (device thickness = 77 μm). Both cases fall far from hemodynamically significant stenosis situations that occur at 50% lumen reduction.^[36]^ It must also be noted that the release in a rigid PMMA block is the worst-case scenario. A more plausible behaviour of the device in-situ would be investigated using mock vessels, with mechanical properties closer to the vasculature allowing for a better device’s apposition. Also, even though the device structure is stent inspired, the device’s primary function is not one of the stents (opening a clogged vessel). Therefore, the expansion at the deployment must be sufficient enough to put the electrodes in contact with the vessel walls, but it will not expand the vessel structure as a stent would do.

**Figure 6.**
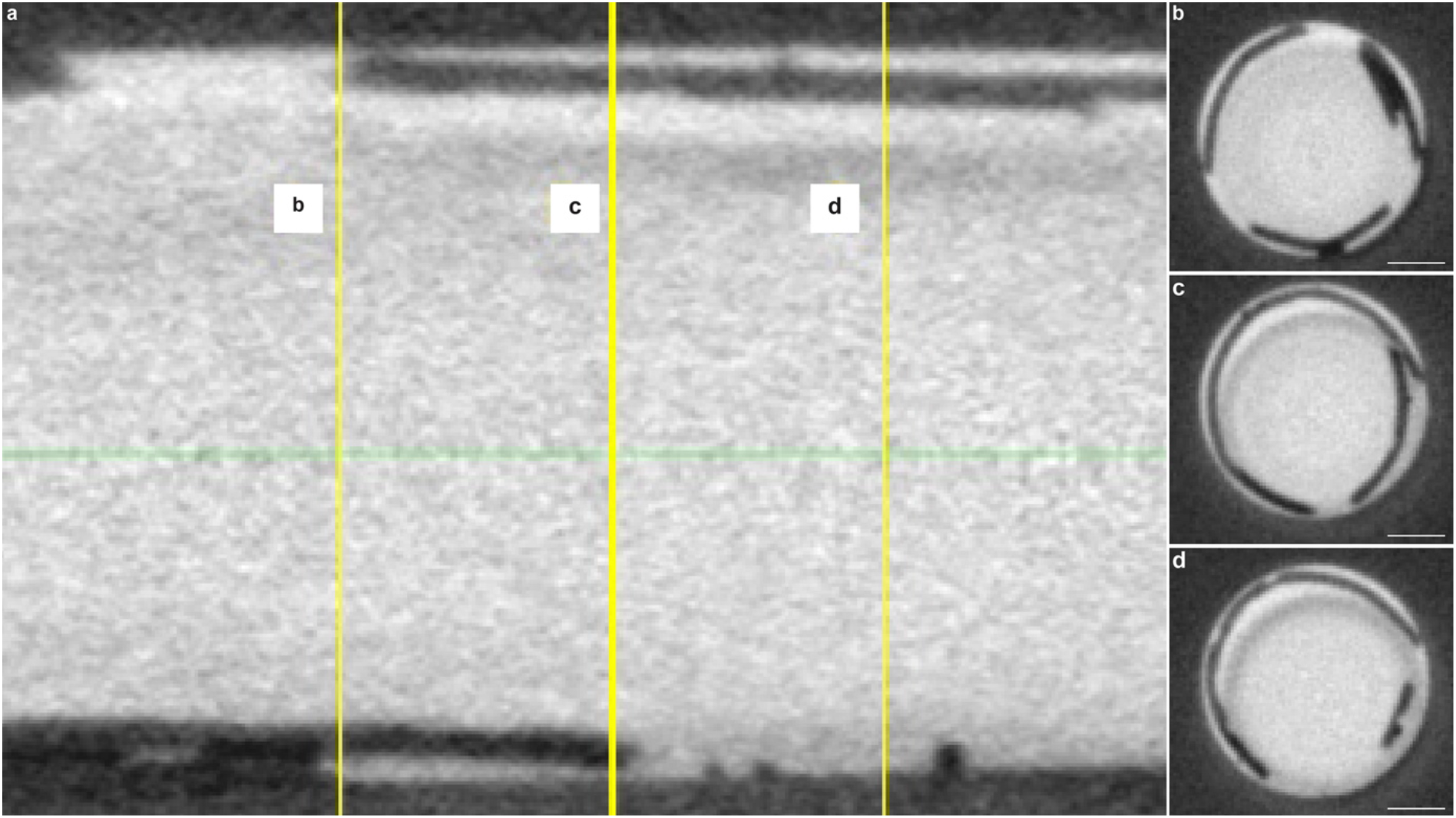
Micro-CT images of the device deployed inside a 2-mm diameter PMMA rigid channel: (a) longitudinal cross-section, (b-d) transversal cross-sections corresponding to the yellow b, c, d, positions in (a). In bright is the contrast agent injected inside the channel. The scale bars in panels (b-d) are 500 μm.

## 3. Conclusion

This work aimed at developing a minimally invasive biodegradable neural interface to target the neural tissue from within a blood vessel.

The device has been successfully fabricated with a simple process employing standard cleanroom techniques, which allows for better electrode’s encapsulation and device’s reproducibility compared to handmade fabrication on metallic stents. Nonetheless, it must be noted that it involved a manual step for the deposition of the PEDOT:PSS-based conductive solution, which could affect the device’s reproducibility. The advantage of the current PEDOT:PSS deposition system inside grooves is the low amount of conductive polymer needed to fill a track (approximately 2 μl). This solution limits substantial waste of material, for example, such as during spin-coating.^[37]^ Patterning PEDOT:PSS using lithography techniques either by lift-off or using a protective layer would be challenging since the compatibility of these processes with synthetic biodegradable polymers such as PCL is limited by solvent’s compatibility or temperatures higher than 50 °C.^[38]^ An exciting solution could involve using a printing system to automatically move a microinjector according to the traces pattern and, at the same time, deposit the conductive polymer homogeneously. Printing of PEDOT:PSS-based solutions is an open research topic, and a standard protocol is not yet defined. So far, mostly ink-jet printing has been used for patterning PEDOT:PSS on solar cells, low-cost in vitro electrophysiology and circuitry design on textiles.^[37–41]^

The device has been successfully navigated into a standard 5-Fr catheter and deployed inside a 2-mm diameter channel built in a rigid PMMA block. After deployment, the device induced a lumen reduction of only 18%, largely below the threshold to observe hemodynamically significant stenosis (50%). However, the validated delivery system uses a guidewire as a support for the device. The two are attached with glue to ensure effective navigation. Once the device is deployed, it would be ideal detaching the two elements. Possible strategies could involve using a temporary glue that would dissolve after deployment or the use of coil embolization wires. The latter is a guidewire with a detachable coil at the end. The detachment is carried out by applying a current to the wire that leads to breaking the terminal coil’s attachment point. If the presented device were attached to the coil, it would be possible to retrieve the guiding wire in this way. Nevertheless, the Stentrode™, built over a stent-retriever consisting of a Nitinol stent attached to a metallic shaft, is currently being validated in a clinical trial with its shaft used as support for the wires and, thus, chronically implanted in the superior sagittal sinus through the transverse sinus.^[3,5]^ Therefore, unless detachable solutions are found, in-vivo validation of the device with the current delivery system can still be considered.

Overall, these results set a good starting point for acute experimentation in-vivo. Future work will also focus on the hemocompatibility of the device.

## 4. Experimental Section/Methods

### Fabrication

Devices were fabricated using a moulding technique. Two master moulds were fabricated: one with protruding pattern of pads, feedlines and electrodes and the other one with engraved parts in correspondence of the feedlines and pads. The moulds were fabricated by SU8 GM1070 coating (1500 rpm, 35 μm) and patterning on 4-inch Si wafers, using standard photolithography (MLA150, Heidelberg Instruments). After hard baking of the moulds, a release layer (poly-styrenesulfonic acid 18 wt% in H_2_O; 561223, Sigma Aldrich) was spin coated on both of them and baked at 145°C for 10 min. Then, poly-ε-caprolactone (MW 50,000; 25090, Polyscience, Inc.) was dissolved in chloroform (C2432, Sigma Aldrich) at 30% w/v and spin coated at 800 rpm on both moulds. A soft bake for 30 min at 75 °C removed residual stresses caused by the spin coating of the viscous solution. Solvent evaporation followed, leaving the wafers under a chemical hood at room temperature for 1 hr. Samples outlines and electrodes openings were laser cut (Optec MM200-USP) and then samples were released by immersion in deionized water. The substrate layer was filled with a solution of PEDOT:PSS (M122 PH1000, Ossila) mixed with 20 wt% ethylene glycol (324558, Sigma Aldrich) and 1 wt% (3-Glycidyloxypropyl)trimethoxylane (440167, Sigma Aldrich), before curing overnight in an oven at 37 °C. After washing the resulting substrate in deionization water to remove uncross-linked PEDOT:PSS monomers, Teflon coated stainless steel wires (114-μm diameter; Z-790500, Science Products GmbH) were connected to the pads using silver paste (G3692 Acheson Silver DAG 1415, PLANO GmbH). The encapsulation layer was then aligned to the substrate and attached to it by quickly melting the two layers at 60 °C.

### Electrochemistry

Electrochemical characterization was performed in a three-electrode configuration with a large area platinum counter electrode and a non-current carrying reference electrode (Ag/AgCl; BASMF2056, Sigma). The electrolytic cell was connected to a potentiostat (Compact Stat, Ivium) and electrochemical measurements were taken at room temperature with each device immersed in phosphate buffered saline at pH 7.2. Impedance spectroscopy was carried out between 1 Hz and 10^5^ Hz using a 50-mV AC voltage. Cyclic voltammetry curves were obtained by sweeping a cyclic potential at 50 mV s^−1^ scan rate between −0.9 V and 0.8 V. For each electrode, the average response over 5 cycles was calculated and the total charge storage capacity was computed from the integration of the respective current curves.

### Electrochemistry under pulsatile flow

A 2-mm diameter channel was fabricated with polydimethylsiloxane (Sylgard 184, Dow Corning) by moulding. The device was inserted in the channel and connected to the tube (Nr.16 MasterFlex L/S PharMed) encased in the rotatory head (7518-10, Masterflex L/S Easy-Load) of the peristaltic pump (Ismatec). The tube and the channel were immersed in PBS (pH 7.2) inside a wide glass beaker placed onto a hotplate (Super-Nuova+, Thermo Scientific) whose temperature probe inside the PBS was set to 37 °C. The pump was set at 500 rpm. Electrochemical measurements of impedance spectroscopy and cyclic voltammetry were performed over time with the device inside the channel, using the beaker as electrolytic cell with Ag/AgCl reference and platinum counter electrodes immersed during measurements.

### Voltage transients’ measurements

Voltage transients were performed in the same three-electrode configuration as electrochemistry. The current pulses were asymmetric (cathodic/anodic ratio = 0.2), charge-balanced and cathodic first. First, electrodes were stabilized with low current pulses (100μA or 10μA, depending on the electrode capacity) at 150 Hz. Stabilization occurred within 6’000 pulses and was followed by higher current stimulations until the maximum cathodal polarization (E_cm_) of the electrode exceeded the water window (−0.9 V - 0.8 V). The E_cm_ was defined as the voltage value measured after 10 μs from the end of the cathodic pulse to avoid the ohmic drop. The maximum injectable current was defined as the maximum current with which the electrodes could be stimulated without their response exceeding the water window limit. The charge injection capacity was calculated by multiplying the maximum injectable current by the pulse width (cathodic phase) and dividing the obtained charge by the electrode geometrical surface area. The long-term stimulation was carried out by stimulating the electrodes of one device at 500 μA cathodic amplitude and 150 Hz up to 100’000 pulses.

### Degradation

Different ratios of PCL:PLGA were mixed and the degradation rates of the relative samples have been investigated. The tested PCL:PLGA ratios were: 100:0, 80:20 and 70:30. The powder mix was dissolved in chloroform (C2432, Sigma-Aldrich) at 45 °C and with magnetic stirring at 200 rpm for 2 hr. For each material combination 3 samples were prepared. The weight of the samples was recorded after fabrication and kept as a reference. Samples were immersed in 10-ml phosphate buffered saline at pH 12 at 37 °C. At each time point samples were taken out of the solution and dried at room temperature under vacuum for 4 hr. Once dry, samples were weighted and, depending on the time point, scanning electron microscopy images were taken.

### Delivery system

The top extremity of device was attached to the end of a push wire (STABILIZER 527-300E, 0.36mm diameter, Cordis) using super glue (Supergel, UHU). The push wire with the device attached was then immersed in deionised water, and the target catheter (5Fr, NavienTM A+ 058, Intracranial Support Catheter, OD 1.7mm) was filled with deionised water. This wetting is used to improve insertion and navigation of the device inside the catheter. The device attached to the push wire was then pushed inside the catheter and navigated until exiting after approximately 1-m of navigation.

### Microtomography imaging

A channel was created in polymethyl methacrylate with a 2-mm diameter. The device was deployed inside the channel as described before. To avoid metallic artefacts generated by the push wire, the attachment site was cut after deployment. To improve the device visibility within the channel, the channel was filled with a contrast agent (Iopamiro 300mg ml^−1^, Bracco) mixed with NaCl solution at 30% v/v. Imaging was performed at 10-μm resolution with a 5-mm field of view (Qunatum CT Lab GX, Rigaku microCT Technology and Perkin Elmer software).

### Scanning electron microscopy

Images were taken with a Schottky field emission scanning electron microscope (SU5000, Hitachi) at 1 kV access voltage and 10% of maximum spot intensity.

### Cytotoxicity test

A test on extract was performed with devices sterilized by UV exposure, with a ratio of the product to extraction vehicle of 3 cm^2^ ml^−1^. The extraction vehicle was Eagle’s minimum essential medium (11090081, Thermo Fisher Scientific) supplemented with 10% foetal bovine serum (10270106, Thermo Fisher Scientific), 1% penicillin-streptomycin (15070063, Thermo Fisher Scientific), 2 mM L-Glutamine (25030081, Thermo Fisher Scientific), and 2.50 μg ml^−1^ Amphotericin B (15290026, Gibco-Thermo Fisher Scientific). The extraction was performed for 24 hr at 37 °C and 5% CO_2_. L929 cells (88102702, Sigma) were plated in a 96 well plate at a sub-confluent density of 7000 cells per well in 100 μl of the same medium. L929 cells were incubated for 24 hr at 37°C and 5% CO_2_. After incubation, the medium was removed from the cells and replaced with the extract (100 μL per well). After another incubation of 24 hr, 50 μL per well of XTT reagent (Cell proliferation kit 11465015001, Sigma) were added and incubated for 4 hr at 37 °C and 5% CO_2_. An aliquot of 100 μl was then transferred from each well into the corresponding wells of a new plate, and the optical density was measured at 450 nm by using a plate reader (FlexStation3, MolecularDevices). Clean medium alone was used as a negative control, whereas medium supplemented with 15% of dimethyl sulfoxide (D2650-5X5ML, Sigma) was used as a positive control. Each condition was tested in triplicates.

### Statistical analysis and graphical representation

Statistical analysis and graphical representation were performed with Prism (Graph Pad). Normality test was performed in each dataset to justify the use of a parametric or non-parametric test. The box plots always extend from the 25^th^ to 75^th^ percentiles. The line in the middle of the box is plotted at the median. The + is the mean. The whiskers go down to the smallest value and up to the largest. In each figure p-values were represented as: * p < 0.05, ** p < 0.01, *** p < 0.001, and **** p < 0.0001.

## Acknowledgements

This work was supported by École polytechnique fédérale de Lausanne and Medtronic.

